# Plant Intron Splicing Efficiency Database (PISE): exploring splicing of ∼1,650,000 introns in Arabidopsis, maize, rice and soybean from ∼57,000 public RNA-seq libraries

**DOI:** 10.1101/2022.06.01.494286

**Authors:** Hong Zhang, Jinbu Jia, Jixian Zhai

## Abstract

Intron retention is the most common form of alternative splicing events in plants and plays a crucial role in plants responding to environmental signals. Despite a large number of RNA-seq libraries from different treatments and genetic mutants stored at public domains, a resource for querying the intron splicing ratio of individual introns is still missing. Here, we established the first-ever large-scale splicing efficiency database to date in any organism. Our database includes more than 57,000 plant public RNA-seq libraries, including 25,283 from Arabidopsis, 17,789 from maize, 10,710 from rice, and 3,974 from soybean, and in total covers more than 1.6 million introns in these four species. In addition, we have manually curated and annotated all the mutant-and treatment-related libraries as well as their matched controls included in our library collection, and added graphics to display intron splicing efficiency across different tissues, developmental stages, and stress-related conditions. The result is a large collection of 3,313 treatment conditions and 3,594 genetic mutants for discovering the differentially regulated splicing efficiency. Our online database can be accessed at https://plantintron.com/.

## INTRODUCTION

Splicing is an essential process during gene expression in eukaryotes and is performed by a large RNA and protein complex called the spliceosome, including U1, U2, U4, U5, and U6 snRNPs and other related proteins (Laloum, *et al*., 2018). Alternative splicing (AS) is a critical process to enrich gene function and plays an important role in response to the environmental signals and genetic mutation during plant development (Staiger and Brown, 2013). Intron retention (IR) is the major event of AS in plants, accounting for more than 40%∼60% of AS events in Arabidopsis (Chaudhary, *et al*., 2019; Reddy, *et al*., 2013), 83% in Oryza sativa (Dong, *et al*., 2018), 56% in maize inbred lines B73 (Mei, *et al*., 2017). And it plays an essential role for plants in gene expression (Jacob and Smith, 2017), responding to environmental signals, such as cold, heat, and drought (Laloum, *et al*., 2018), and regulating splicing efficiency (Chong, *et al*., 2019).

In previous studies, researchers have preferred to focus on one or a limited number of genetic or environmental stresses to investigate the effects of AS or IR, such as the histone methyltransferase SET DOMAIN GROUP PROTEIN 725 (SDG725) mediating position-specific IR in rice (Wei, *et al*., 2018), the protein arginine methyltransferases 5 (PRMT5) involved in the pre-mRNA splicing through Sm-like4 (LSM4) methylation in Arabidopsis, which enhance the high slat tolerance (Zhang, *et al*., 2011), and the phytohormone abscisic acid (ABA) signal pathway is an important pattern to response abiotic stress, and the last intron of *HAB1* would be retained without ABA and *RBM25* (Cheng, *et al*., 2017; Wang, *et al*., 2015; Zhan, *et al*., 2015). Thus, there are large-scale raw data published in the public database. It’s a hefty charge for most researchers to reanalyze all public RNA-seq data which are limited by computer resources or bioinformatics technology. Although there are also some resources for studying AS of plants, such as the PastDB which used 516 public libraries to analyze gene expression levels and AS atlas in Arabidopsis (Martin, *et al*., 2021), ASIP (Wang and Brendel, 2006) and ERISdb (Szczesniak, *et al*., 2013) focus on the characterization of AS events. There is no comprehensive database to investigate intron splicing efficiency by analyzing all public RNA-seq data of plants, and no resource for users to quickly verify whether genic mutations or treatment conditions affect intron splicing and whether some differences exist in different tissues and developmental stages.

Our previous studies collected over 20,000 Arabidopsis RNA-seq public libraries (Zhang, *et al*., 2020) and 4,5000 RNA-seq libraries for five plant species (Yu, *et al*., 2022), which supported an important resource for investigating splicing efficiency. Furthermore, we established the first-ever large-scale splicing efficiency database in four organisms to date in this work (PISE: https://plantintron.com/), which can reveal the intron splicing efficiency of selected gene and intron in multiple formats with tables and figures. Besides, our website includes a web-based IGV browser to display the bam files of all ∼57,000 libraries stored on our server. Users can easily choose the libraries for display by searching library ID, project ID, or gene ID in our collection.

## RESULTS

### Building a comprehensive database for plant intron splicing efficiency

Our database collected more than 57,000 public Next-generation sequencing (NGS) RNA-seq libraries and used the given pipeline for each plant to analyze intron splicing efficiency (IR ratio) (Figure 1A). In addition, we have manually curated and annotated all the mutant-and treatment-related libraries as well as their matched controls included in our library collection, and added graphics to show intron splicing efficiency across different tissues, developmental stages, and stress-related conditions. As a result, we obtained a large collection of 3,313 treatment-and 3,594 genetic mutant-related groups to discover the differentially regulated splicing efficiency. Finally, we developed a friendly, easy accessing, large-scale database (PISE) to search intron splicing efficiency (Figure 1B), which included over 25,000 libraries for Arabidopsis, 3,000 libraries for soybean, 17,000 libraries for maize, and 10,000 libraries for rice (Figure 1C).

**Figure 1.**
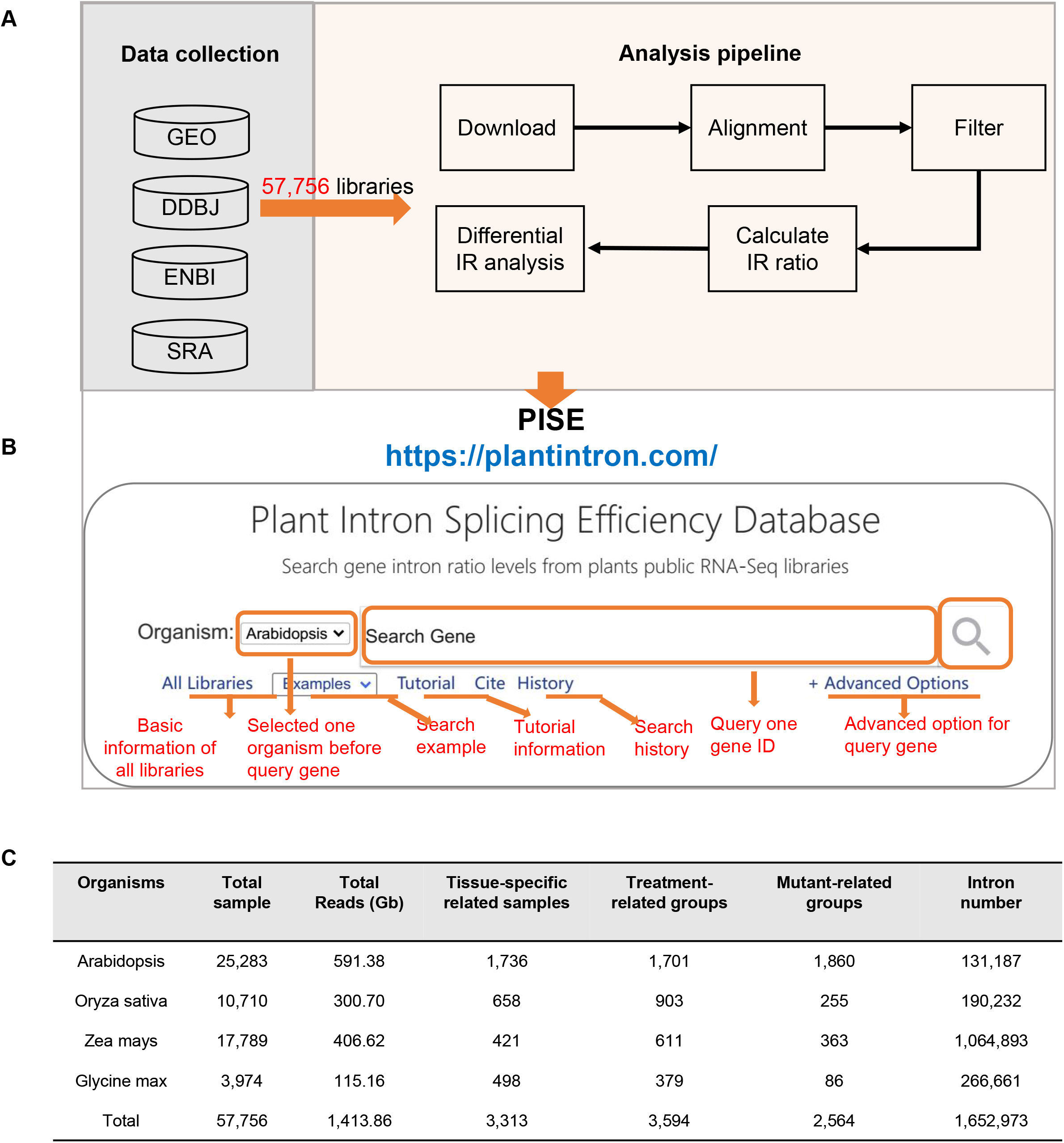
Overview of Plant Intron-Splicing Efficiency Database (PISE). **A**. The pipeline of data collection and process. **B**. The home page of PISE. **C**. Statistics on the number of public RNA-seq libraries and introns used in PISE.

### Overview of selected gene’s splicing information

To overview all intron splicing efficiencies of the selected gene, we show the IR ratio among all libraries of selected organisms in the “Information” page, including a box plot of all intron retained levels of selected transcript (Figure 2A) and a basic statistical table (Figure 2B), such as IR ratio in all libraries for each intron, the median, mean, the third quartile (Q3) value for each intron. To facilitate access to each intron to scan the detailed information, we have added a function event for each point in the plot area to show the library name and a function for box clicking to show the IR ratio value of each library. Users can query one intron by clicking the intron ID in the table to get more information about the selected intron. Besides, we have changed the function of the legend so that clicking on the legend will also display information about the chosen intron. The result information of chosen intron will be displayed on the “Data Table” and “Data Plot” pages.

**Figure 2.**
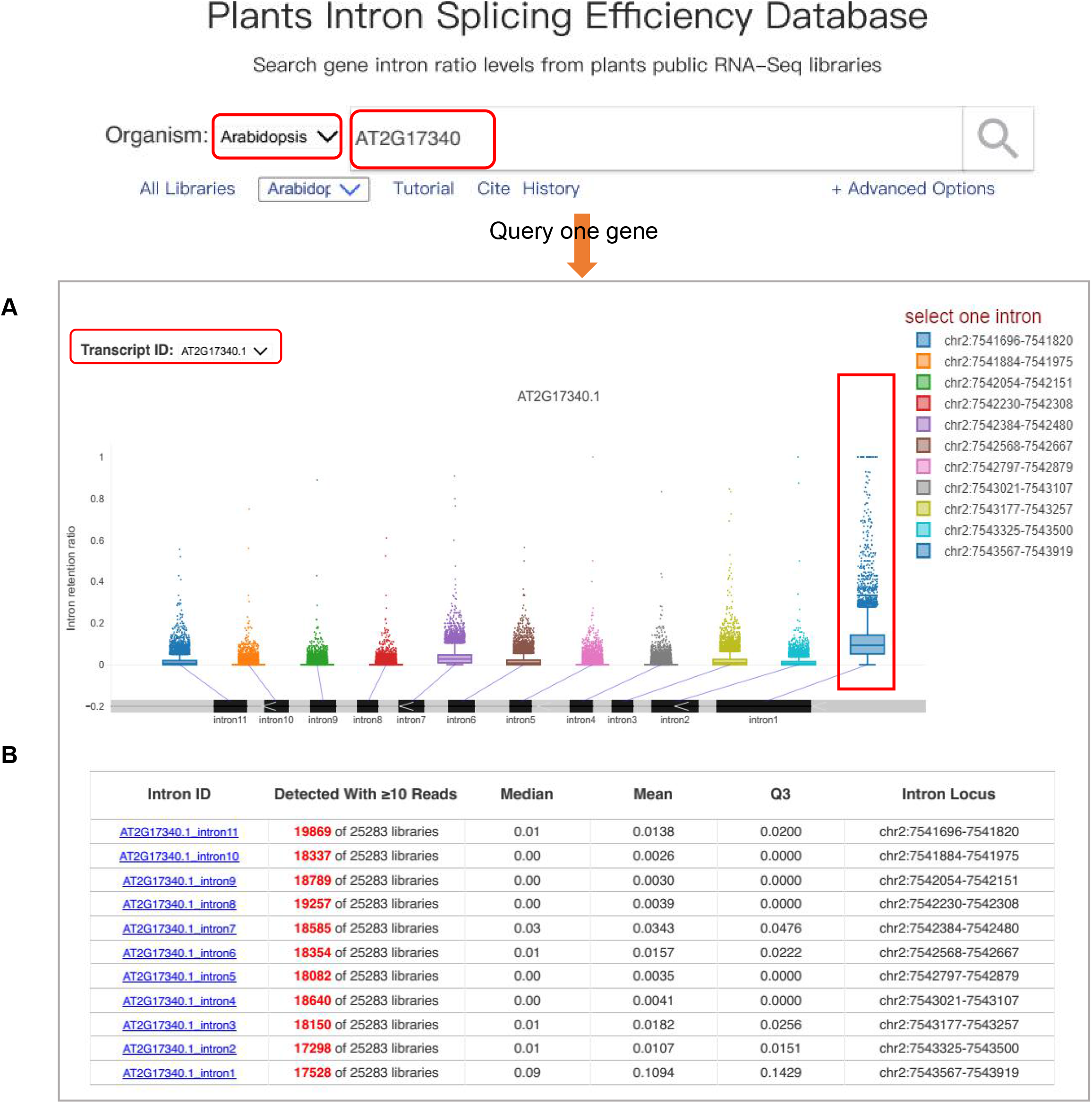
The result after querying one gene. **A**. The box plot of intron retention ratio across AT2G17340.1’s all intron in Arabidopsis. Users can find other transcripts’ intron splicing levels in all public RNA-seq libraries by changing the ‘TranscriptID’ and can access the plot data by clicking the box. **B**. The drawn intron basic information. The legend in A and” Intron ID” in B both supported the function of searching one intron by clicking intron ID.

### Multi-format of the queried intron

*Data Table*. After selecting one intron, the “Data Table” page will show the detailed information about IR ratio in all libraries, including sample name, title, IR ratio, reads covered within intron range, all reads within the intron flank range, ecotype/cultivar, genotype, tissue, project ID, treatment, release date (Figure 3A). The load table function is supported by canvas-datagrid, which provides a right-clicking filtering function for each column. To facilitate filtering, we have also added the filter button above the table to support filtering by the reads number of the selected intron, project, tissue, ecotype, and genotype (Figure 3A). Moreover, we supported the table download function to download the details of the selected intron to perform data mining. For example, the previous study has reported that the first intron of AT2G17340.1 is largely retained on the *atprmt5* mutant (Deng, *et al*., 2016), and this result can be discernible in our database (black point in Figure 3C). In addition, we found that some splicing-related factor mutants or environmental signal-related libraries also have high retained levels on this intron, such as *sac3a-3 (YEAST SAC3 HOMOLOG A)* and *prp4a-1(pre-mRNA processing 4 KINASE A)* mutant, heat and cold stress (Figure 3C). There are some other mutant libraries, such as *rrc1-3(REDUCED RED-LIGHT RESPONSES IN CRY1CRY2 BACKGROUND 1)*, that may be associated with affecting the splicing efficiency of this intron and can be tested via some experiments.

**Figure 3.**
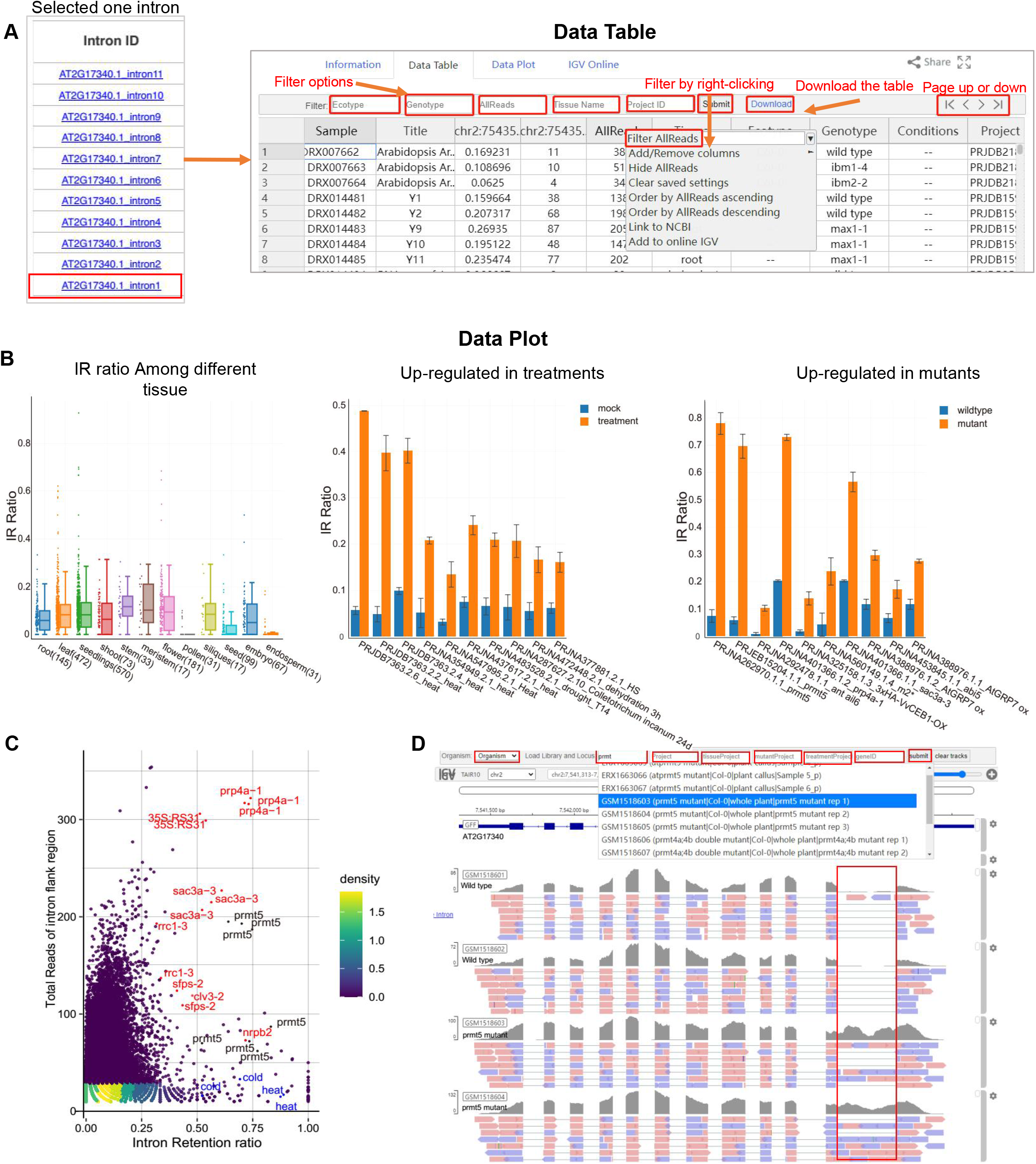
Result of the selected intron. **A**. A table for the selected intron splicing efficiency on all public libraries, including library title, project, genotype, ecotype, conditions, the retention ratio, and flanking exon reads. **B**. Multi-plot of the selected intron (AT2G17340.1#intron1), the left panel is the retention ratio among different tissue, the middle panel is the increased retention level in the top 10 treatment conditions, and the right panel is the up-regulated retention level in the top 10 mutants. **C**. Intron retention levels of AT2G17340.1#intron1 in all Arabidopsis libraries. *prmt5* mutant, which detected IR event of this intron in a previous study, was marked black, other mutants with high IR ratios were labeled red, and abiotic stress-related samples with high IR ratio were marked in blue. **D**. Coverage of AT2G17340 in the selected libraries.

#### Data Plot

To better display the selected intron for the user, “Data Plot” supports multi-format plots to show the intron retained levels, including IR ratio across different tissue, development stages, biotic and abiotic stresses, as well as the up-or down-regulated between treatment and matched mock, between mutants and matched wild-type libraries (Figure 3B). In addition, box plots have been used to show IR ratio among all related samples of the tissue-specificity, development stage, biotic, and abiotic stress, and bar plots have been used to display the top 10 groups showing differential intron splicing efficiency among related mutants and treatments. The figure and corresponding data download function also be supported in the plot area, which allows users to download relevant data by clicking the related button. Taking the first intron of AT2G17340.1 as an example, by the result of differential intron retention analysis, we can find that the intron retained level has been up-regulated under some mutants and treatment conditions, such as *prmt5, ant/ail6, prp4a-1, sac3a-3* mutants, heat and drought treatments (Figure 3B). This coincides with previously reported retention of this intron in *prmt5* (Deng, *et al*., 2016) and directly overlaps with what was found in “Data Table”.

### IGV browser

The “IGV Online” page shows the functionality of the genomics viewer (Robinson, *et al*., 2017). Users can easily view the covered reads for selected genes in all public libraries of the chosen organism without any processing of the raw data. For example, we showed the covered reads of *atprmt5* mutant libraries and matched controls of AT2G17340 in Arabidopsis (Figure 3D) and found that the coverage of the first intron was significantly higher in the mutant libraries. This result is consistent with findings in Figure 3B and Figure 3C, visually demonstrating the high reliability of our analysis. Furthermore, each button supports an auto-complete function that allows users to search relevant libraries or genes by keyword or ID.

## DISCUSSION

We support a free comprehensive database for quickly scanning plant intron splicing efficiency. Users can easily access splicing information for the interesting intron in public data without any data processing, including how many reads support intron retention, what treatments will up-or down-regulated the splicing efficiency, and whether the intron is tissue and development stages specifically regulated. Even though there are some resources for investigating plant AS, they focused on the splicing site features (Szczesniak, *et al*., 2013; Wang and Brendel, 2006) or only used 516 public libraries to study the splicing atlas (Martin, *et al*., 2021). Our database used the larger-scale public data and showed the intron splicing efficiency using data tables and figures in multiple formats of four plants, making it more user-friendly to get valid or interesting information.

This way, most researchers with limited informatics background could benefit from our website using a “data-driven, hypothesis-generating” approach without performing sophisticated informatics analysis, and come up with testable experimental hypotheses based on novel information extracted from such a comprehensive database. For example, the first intron of AT2G17340.1 was reported to be retained in Arabidopsis *prmt5* mutant (Deng, *et al*., 2016), and our database not only confirm this result with *prmt5* mutant libraries but also discovered many other potential regulators for this intron from the mutant analysis (Figure 3B-C). In a word, PISE provides a concise, friendly, “Google-style” search function to query gene intron splicing efficiency, allowing users easy to obtain their interesting gene or intron’s splicing levels under different tissue, development stage, stresses, treatments, and genetic mutants.

## MATERIAL AND METHODS

### Data collection and process

We collected the public RNA-seq data from Gene Expression Omnibus (GEO), the Sequence Read Archive (SRA), the European Nucleotide Archive (ENA), and the DNA Data Bank of Japan databases (DDBJ) for Arabidopsis, rice, maize, and soybean until 2020, then manually went through all libraries to complete the basic information and filter some libraries which may be smRNA or lncRNA. The raw data were aligned to the reference genome by HISAT2 (Kim, *et al*., 2015), and the TAIR10, Williams 82, B73 v4, and MSU 7 genomes were used as references for Arabidopsis, soybean, maize, and rice, respectively. Then, libraries with low mapped reads (unique mapped reads < 1M) were discarded. Finally, we used over 57,000 libraries (Figure 1C) to perform the downstream analysis. Only genes with at least one intron were used for analysis, and the intron splicing efficiency (IR ratio) was calculated by a python script from Jia (Jia, *et al*., 2020). All intron information was extracted from published annotation files, including Araport11 for Arabidopsis, B73 v4 for maize, MSU 7 for rice, and Williams 82 a2 for soybean.

### Differential intron-splicing efficiency analysis

After filtering, we manually curated and annotated all the mutant-and treatment-related libraries as well as their matched controls included in our library collection and added graphics to display intron splicing efficiency across different tissues, developmental stages, and stress-related conditions for Arabidopsis, maize, rice, and soybean. The result is a large collection of 3,313 treatment conditions and 3,594 genetic mutants for discovering the differentially regulated splicing efficiency (Figure 1C), which at least two libraries with more than 1M mapped reads for the control and treatment/mutant groups, respectively. The generalized linear model in DESeq2 (Love, *et al*., 2014) was used to calculate the fold-change and p-value for each intron as well as be used in the IRFinder test (Middleton, *et al*., 2017), and the Benjamini Hochberg method was used to adjust the p-value (p-adj). And the fold change and p-adj were filtered for each intron before being plotted on the “Data Plot” page.

### PISE website and database

MySQL and LMDB were used as storage units to hold all relevant data tables for querying genes or libraries, including the library’s basic information, gene information, and corresponding values of intron splicing efficiency. Php, HTML, and JavaScript were used to build a web framework that connected PC and server. To better display the data, we upgraded the function of Plotly.js to plot all figures and support downloading and displaying the corresponding data, and added a filter button in canvas-datagrid.js to load the data table. We used the parameters “p-adj<0.05 and mean of control or experiment group IR ratio ≥ 0.1” to filter all related groups for mutant-and treatment-related plots and plot the retained groups’ libraries. To conveniently view one gene’s coverage of selected libraries online, the online genomic browser was supported using igv.js (Robinson, *et al*., 2017), which supports library ID, project ID, treatment, mutant, tissue groups ID, and keywords to search interested library.

### Compliance and ethics

The author(s) declare that they have no conflict of interest.

## Acknowledgement

We thank all the research groups that contributed RNA-seq data to the community, and we apologize for not being able to cite all the related papers in the main text due to limited space. References for all libraries we used are listed in PISE ‘All Libraries’. Computation was supported by Center for Computational Science and Engineering at Southern University of Science and Technology. The group of J.Z. is supported by the National Key R&D Program of China Grant (2019YFA0903903); the Program for Guangdong Introducing Innovative and Entrepreneurial Teams (2016ZT06S172); the Shenzhen Sci-Tech Fund (KYTDPT20181011104005); the Key Laboratory of Molecular Design for Plant Cell Factory of Guangdong Higher Education Institutes (2019KSYS006); and the Stable Support Plan Program of Shenzhen Natural Science Fund Grant (20200925153345004). J.J. is supported by the National Natural Science Foundation of China (32100444); and the Shenzhen Fundamental Research Program (JCYJ20210324105202007).

